# Accelerated simulation of evolutionary trajectories in origin–fixation models

**DOI:** 10.1101/087700

**Authors:** Ashley I. Teufel, Claus O. Wilke

## Abstract

We present an accelerated algorithm to forward-simulate origin--fixation models. Our algorithm requires on average only about two fitness evaluations per fixed mutation, whereas traditional algorithms require, per one fixed mutation, a number of fitness evaluations on the order of the effective population size Ne. Our accelerated algorithm yields the exact same steady state as the original algorithm but produces a different order of fixed mutations. By comparing several relevant evolutionary metrics, such as the distribution of fixed selection coefficients and the probability of reversion, we find that the two algorithms behave equivalently in many respects. However, the accelerated algorithm yields less variance in fixed selection coefficients. Notably, we are able to recover the expected amount of variance by rescaling population size, and we find a linear relationship between the rescaled population size and the population size used by the original algorithm. Considering the widespread usage of origin--fixation simulations across many areas of evolutionary biology, we introduce our accelerated algorithm as a useful tool for increasing the computational complexity of fitness functions without sacrificing much in terms of accuracy of the evolutionary simulation.

## 1. Introduction

A key goal in the field of molecular evolution is to understand the processes by which proteins acquire substitutions and diverge from one another. The phenotypic effect a mutation has and whether it ultimately becomes fixed or lost in a population both depend in part on the biochemical and biophysical effects the mutation has on protein structure and function [1–5]. By studying the physio-chemical forces which underlie the substitution process we can begin to untangle the mechanisms which have given rise to organismal diversity, shedding light on linage-specific functional divergence [6–9].

While considerable progress has been made in characterizing the substitution process through mathematical modeling and from studying extant sequences, further understanding can be gleaned from observing evolution directly via simulation. However, simulating fitness landscapes that explicitly and accurately model the effects of individual mutations on protein structure and function can be computationally burdensome. Traditionally, this problem has been addressed by using coarse-grained protein models, such as lattice or bead models, due to their computational tractability [3,10]. Here, we take a different approach to solving the problem of computational tractability. Rather than relying on highly simplified fitness landscapes, we propose a more efficient algorithmic strategy for simulating evolution.

Our approach is based on the origin–fixation model, which applies in the limit of strong selection and weak mutation [11]. In this model, populations are assumed to be monomorphic and represented by a single genotype. Mutations are introduced sequentially (origin) and either accepted or rejected (fixation). This model has been applied in a number of different context [12–18] to simulate evolution over various fitness landscapes, and it is of course much more computationally efficient than simulating all individuals in a population. However, even origin–fixation simulations can be computationally expensive. These simulations require fitness calculations for a large number of genotypes, and the majority of these genotypes will be rejected and not influence the evolutionary path taken. For fitness landscapes where the evaluation of a single genotype is computationally expensive, for example when fitness is based on detailed, atomistic models of protein structure and dynamics, it is desirable to minimize the number of these evaluations. The fewer fitness evaluations we have to perform per accepted mutation, the more computational time we can expend on each individual fitness calculation, and hence the more realistic a model we can simulate.

In the context of protein evolution, recent work has argued, based on first principles, that most fixed mutations have to be neutral or nearly neutral [19,20]. In this scenario, we expect that for one fixed mutation, we have to evaluate and discard a number of mutations on the order of the effective population size *N*_*e*_. Discarding that many mutations after evaluating their fitness is a highly inefficient way of simulating evolution, and any reduction in the number of fitness evaluations required per fixed mutation will speed up the simulation. Here, we demonstrate that by using a modified acceptance criterion for fixed mutations, we can reduce the number of required fitness evaluations to approximately two per one fixed mutation, a speed-up on the order of *N*_*e*_. Our algorithm, which uses a scaling idea originally introduced in the context of Metropolis–Hastings sampling [21,22], is guaranteed to yield the same evolutionary steady state as the original origin–fixation model. Further, we demonstrate through extensive simulations with toy models and with an all-atom, energy-based protein model that the key difference of the accelerated algorithm is reduced variance in fixed selection coefficients. The magnitude of reduced variance is small, on the order of the difference between a Wright-Fisher model and a Moran model, and it can be compensated by choosing a slightly smaller effect population size *N*_*e*_.

All simulation models make assumptions about how the underlying processes operate, and the relative validity of these assumptions hinges on the specific context in which a model is used. Our accelerated algorithm presents a useful way to rapidly simulate protein evolution in the context of a detailed, atomistic model of protein structure. However, our algorithm does perturb the evolutionary process and produces data that may not be appropriate for some types of analysis. Ultimately, the balancing point between more detailed descriptions of individual fitness and the accuracy of an evolutionary model depends on the questions being posed of the data. We present our algorithm as an option along this continuum, favoring a rigorous treatment of individual fitness over evolutionary precision.

## 2. Results

### Evolutionary simulations using Metropolis sampling

Our accelerated algorithm shares the same basic structure as a number of origin–fixation based simulations [11–15]. The simulation is initialized with a monomorphic population represented by a single genotype *i*. At each iteration a novel mutation is tested and is either accepted and becomes the represented genotype or rejected and the current genotype remains. The difference between our algorithm and the original is based on the probability of accepting a mutation. To illustrate how the two acceptance criteria differ we first outline the theory behind the original algorithm and then explain our modification to it.

Consider a population of *N*_*e*_ individuals evolving in discrete time-steps according to Wright-Fisher sampling [23]. Assume the overall mutation rate *μ* is sufficiently low such that *μN*_*e*_ ≪ 1. We denote the fitness of genotype *i* by *f*_*i*_ and the mutation rate from *i* to *j* by *μ*_*ji*_. We assume a symmetric mutation matrix, *μ*_*ij*_ = *μ*_*ji*_. Under these assumptions, the evolutionary process can be described as follows [24]. First, the fixation probability for a mutation from *i* to *j* can be written as

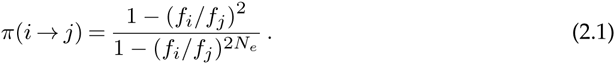

Next, let *W*_*ji*_ define a Markov process where the transition probabilities from state *i* to state *j* are given by

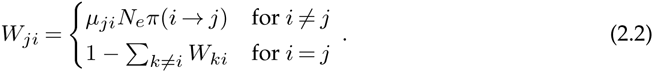

This Markov process describes the origin–fixation model. The term *μ*_*ji*_*N*_*e*_ provides the rate at which mutations from *i* to *j* are generated in a population of size *N*_*e*_, and the term π(*i* → *j*) provides the probability that one of these mutations goes to fixation.

It can be shown [24] that the Markov process *W*_*ji*_ has stationary frequencies

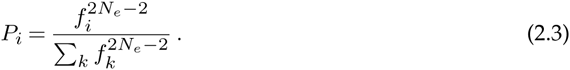

These frequencies satisfy the detailed balance equation

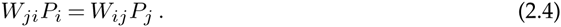

To simulate an origin–fixation model, we start with a genotype *i* and generate a new mutation according to the mutation matrix *μ*_*ij*_. We then accept or reject the mutation according to the fixation probability, as given by Eq. (2.1) [11]. Thus, we can think of the fixation probability as an acceptance criterion for a proposed mutation. If a fitness landscape has a large proportion of neutral or nearly-neutral genotypes, on average on the order of *N*_*e*_ mutations need to be evaluated before one is accepted.

Our modification to this algorithm is based on the fact that *W*_*ij*_ satisfies the detailed balance condition Eq. (2.4). Because of this condition, we can rescale *W*_*ij*_ with any symmetric matrix and retain the same steady-state frequencies. Let 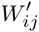 denote a rescaling of *W*_*ij*_ for all non-zero paths in the matrix, such that

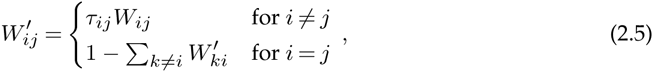

where *τ*_*i,j*_

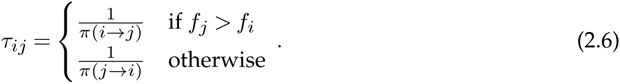

Here *τ*_*ij*_ is defined such that we rescale transitions along the (*I*, *j*) edge with the inverse of the probability of fixing the higher-fitness genotype when starting out at the lower-fitness genotype.

By comparing 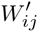 with *W*_*ij*_, we see that we can incorporate *τ*_*ij*_ into a modified fixation probability,

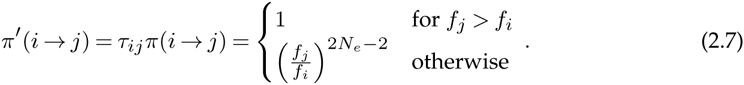

If we interpret this modified fixation probability as an acceptance criterion in an origin–fixation model, we see that, under this new acceptance criterion, we always accept mutations that increase fitness. Mutations that decrease fitness are exponentially unlikely to be accepted.

Notably, Eq. (2.7) looks exactly like the Metropolis-Hastings acceptance criterion widely employed in MCMC sampling [21,22], and indeed, our rescaling approach is equivalent to and was directly motivated by the original work of Metropolis *et al*. [21]. The advantage of Eq. (2.7) over Eq. (2.1) is that we now accept every neutral or beneficial mutation, hence the simulation becomes much more efficient. However, this advantage comes at a cost. Even though the steady state of the simulated Markov process remains unchanged, by construction, the order in which states are visited has changed. Since we use a different rescaling constant along each mutational path, the relative likelihood with which specific paths are chosen has changed between the original and the accelerated process. In particular, in the accelerated process we have much more rapid accumulation of neutral or nearly neutral mutations relative to the rate at which highly beneficial mutations accumulate.

To evaluate the effect of this distortion we performed a set of simulations using a simple three-state fitness landscape. The three states are denoted as A, B, and C, and states A and B are assigned the same fitness (1.0) while state C has a higher fitness (1.25) (Fig 1A). The transition probabilities between states are shown in Fig 1B for the regular acceptance criterion Eq. (2.1) and in Fig 1C for the accelerated acceptance criterion Eq. (2.7). In both cases, we used a relatively small *N*_*e*_ = 10, to ensure the process regularly moves from the higher-fitness state to one of the lower-fitness states.

**Figure 1.**
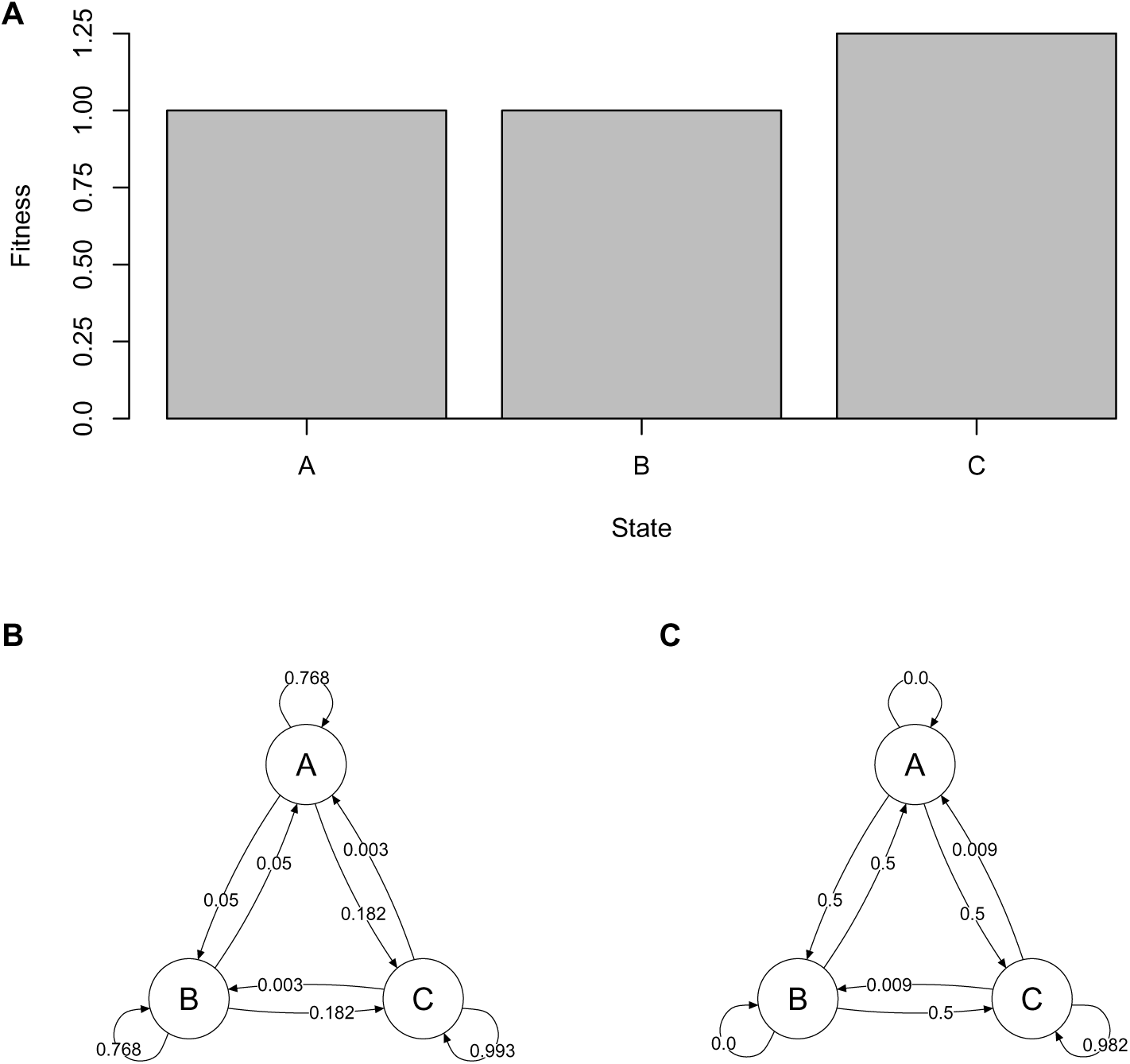
Comparison of original and accelerated sampling algorithm on a simple three-state fitness landscape. (A) States A and B have equal fitness, set at 1.0, and state C has a 25% higher fitness of 1.25. (B) Directed graph showing the transition probabilities between the three states, calculated using the original fixation probability Eq. (2.1) and assuming *N*_*e*_ = 10. (C) Directed graph showing the transition probabilities between the three states, calculated using the accelerated fixation probability Eq. (2.7) and again assuming *N*_*e*_ = 10.

We simulated origin–fixation models over the three-state landscape using both acceptance criteria, and we found that the mean time spent in any state did not differ significantly between the two algorithms (Fig 2A, Table S1), as expected due to the construction of the accelerated algorithm. Interestingly, the variance in the amount of time spent in any state did differ (Table S1). Fig 2B shows that the fraction of types of transitions that occurred (e.g. A to B or B to A) differed between the two algorithms as well (Table S2). Combined, these results suggest that the two algorithms produce distinct patterns in the order states are sampled and in the frequency of transition types. We also inspected the distributions of selection coefficients produced by the two algorithms (Fig S1). Although the distributions appear similar, a Kolmogorov-Smirnov (KS) test revealed that they are not equivalent (*p* = 5.0 × 10 ^−7^).

**Figure 2.**
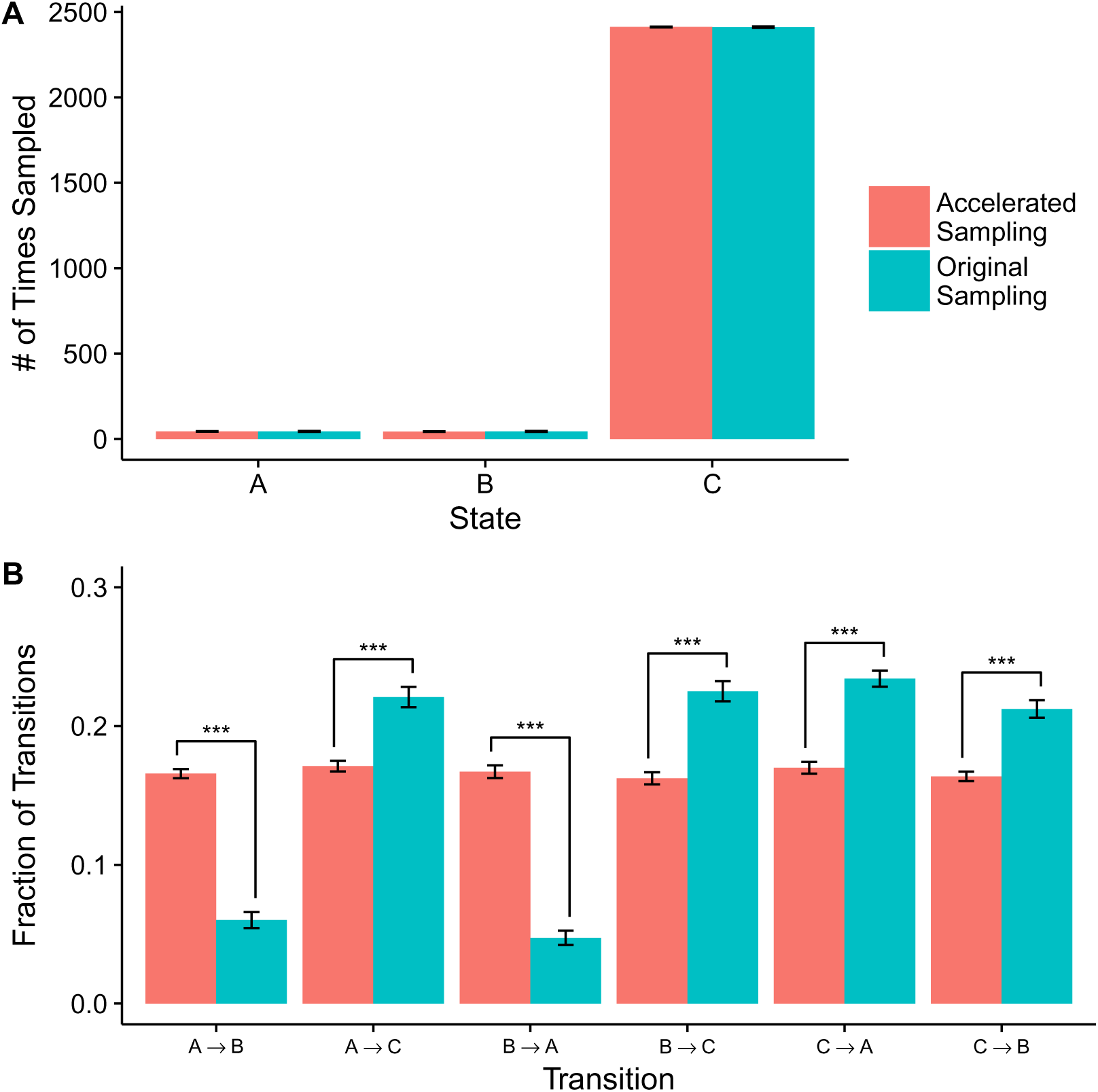
Comparison of the frequencies with which states are visited and with which specific transitions occur in the original and the accelerated sampling algorithm, using the three-state fitness landscape of Fig. 1 and *N*_*e*_ = 10. (A) The mean number of times a state is visited does not differ significantly between the two algorithms (t tests, all *p* > 0:2, Table S1). However, the variance in the number of times a state is visited does differ (Table S1). (B) The mean fractions of specific transitions between states differ between the two algorithms (all *p* < 0:001, indicated by three stars, Table S2).

To assess whether the differences in results generated by the two algorithms were due to the modified acceptance criterion and not some underlying feature of the particular fitness landscape considered, we repeated this experiment on a different three-state fitness landscape. We found that these additional simulations resulted in similar dynamics to those obtained on the initial landscape (Figs S2, S3, and S4, Tables S1 and S2).

### Application to protein evolution

To evaluate the performance of our accelerated algorithm in a more realistic context, we constructed simulations of protein evolution using a computationally costly all-atom model of protein structure. The simulation was initialized with a small protein as the resident genotype. At each iteration a mutation to a non-resident amino acid was introduced to a random location and the structure was locally repacked 10Å around the mutation. The stability (*ΔG*) of the mutant was evaluated with Rosetta’s all-atom score function [25,26]. To convert protein stability into fitness, we used a soft-threshold model. This model assumes that the protein’s fitness is given by the fraction of proteins in the ground state in thermodynamic equilibrium [27–29]. This assumption results in a sigmoidal fitness function (specifically, the Fermi function), where very stable proteins have a fitness of one and fitness declines as stability passes through a threshold value. We calculate the fitness of protein *i* as

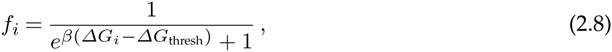

where *β* is the inverse temperature, *ΔG*_*i*_ is the stability of protein *i*, and *ΔG*_thresh_ is the stability threshold at which the protein has lost 50% of its activity.

To simulate the accelerated algorithm, it is convenient to log-transform fitness,

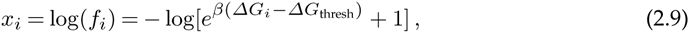

so that the fixation probability Eq. (2.7) can be expressed as (assuming *N*_*e*_ − 1 ≈*N*_*e*_)

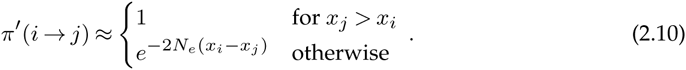

As in Eq. (2.7), the transitions from a lower fitness to a higher fitness always happen, while transitions from a higher fitness to a lower fitness are exponentially suppressed in *N*_*e*_ and in the difference in log-fitness *x*_*i*_ − *x*_*j*_.

The soft-threshold fitness model of protein evolution has been studied previously [28,30], and the relationships it produces between protein stability, population size, and inverse temperature *β* are well understood. As effective population size *N*_*e*_ increases, increasingly smaller fitness differences become visible to selection, and this results in an increase in protein stability in the steady state. The inverse temperature *β* determines how hard or soft the fitness threshold is. For very large values of *β*, the fitness function turns into a hard cutoff, producing a fitness of 0 for *ΔG*_*i*_ > *ΔG*_thresh_ and a fitness of 1 for *ΔG*_i_ < *ΔG*_thresh_. For smaller *β*, the fitness function becomes increasingly shallow, and this more shallow fitness function allows for a larger range of *ΔG* values to be explored at the point of mutation–selection balance. This effect ultimately increases the amount of variance in stability sampled over the course of the simulation.

To compare the original and accelerated algorithm for this protein model, we initially simulated evolutionary trajectories in triplicate under both algorithms and recorded the first 500 accumulated substitutions. In these simulations, we set *N*_*e*_ = 100, *β* = 1, and *ΔG*_thresh_ = *ΔG*_init_/2, where *ΔG*_init_ is the initial stability of the protein. (For the protein used in these simulations, *ΔG*_thresh_ = −267, measured in arbitrary units). Comparing both algorithms, we saw that protein stability *ΔG* equilibrated after about 200 substitutions, and subsequent fluctuations in stability were comparable across algorithms (Fig 3A). The equilibrium stability *ΔG* was close to the threshold value. Notably, the original algorithm exhibited a greater amount of variation in the *ΔΔG* of fixed mutations than did the accelerated algorithm (Fig 3B).

**Figure 3.**
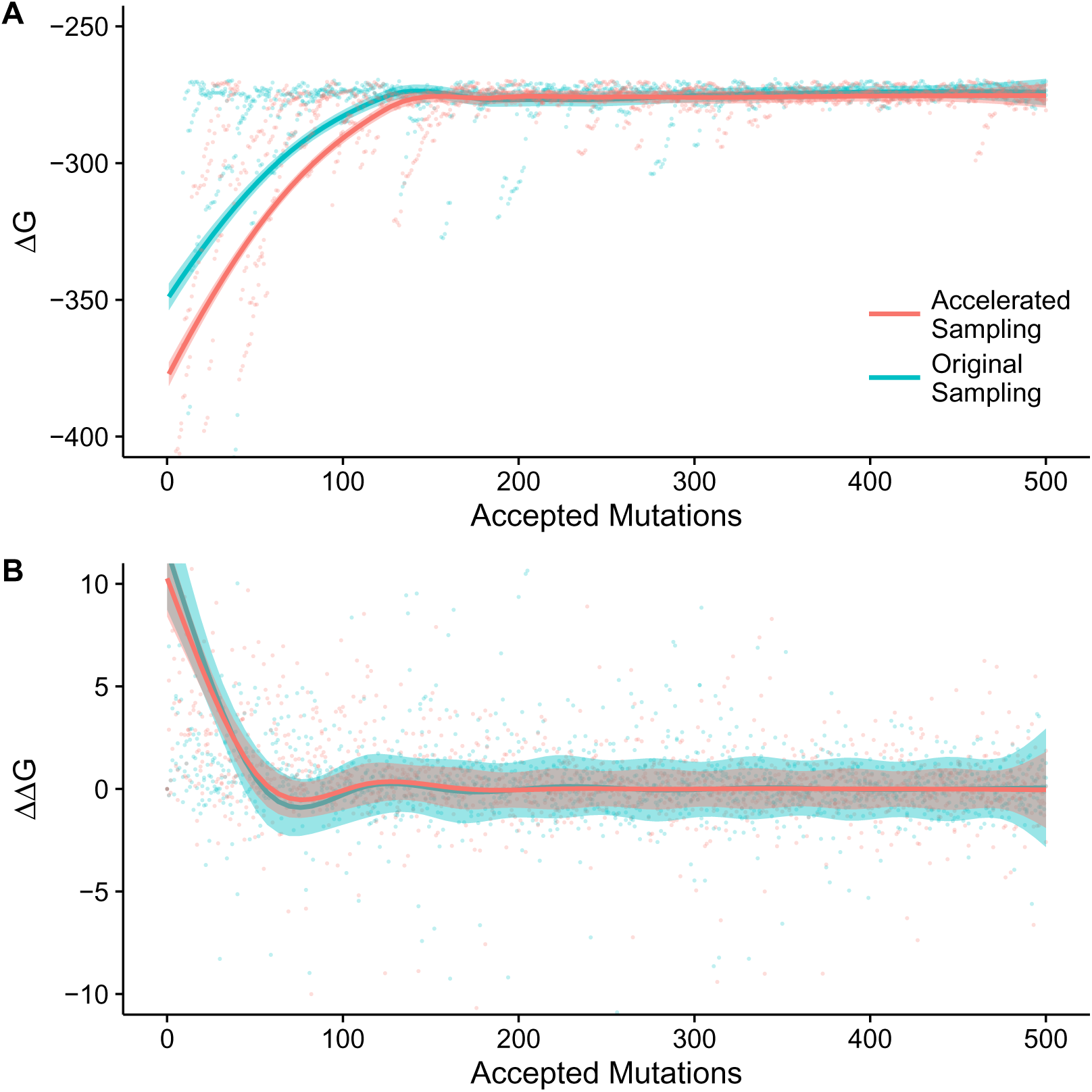
Stability *ΔG* (A) and change in stability *ΔΔG* caused by fixed mutations (B) for the first 500 accepted mutations when simulating the evolution of a small protein under a soft-threshold fitness model. Simulations parameters were *N*_*e*_ = 100, *β* = 1, and *ΔG*_thresh_ = −267. Points represent individual measurements of *ΔG* or *ΔΔG*, pooled over replicate runs. Lines represent a local smoothing of the data, and shaded areas represent the standard error around the smoothed estimate.

The average number of proposed mutations necessary for 500 substitutions was 141,035 using the original algorithm and 1,299 using the accelerated algorithm. The mean run time of the accelerated algorithm was 2.55 minutes and the original algorithm required, on average, 2.73 hours. Thus, we observed a speed-up of approximately 64 fold at a simulated population size of *N*_*e*_ = 100.

We next compared the equilibrium behavior of the original and accelerated algorithms, running simulations for 1,500 accumulated substitutions and discarding the first 1,000 substitutions as burn-in. To examine whether the original and accelerated algorithms produced comparable patterns of divergence, we compared the distributions of the numbers of substitutions at individual sites. We found that both distributions were highly similar (Fig S5A, KS test; *p* = 1). We also compared the specific numbers of substitutions at individual sites. While these numbers are noisy, due to limited sampling, substitution numbers at individual sites were significantly correlated (Fig S5B, Pearson’s *R* = 0.603, *p* < 2.2 × 10 ^−16^). Moreover, the site-specific variation among algorithms was comparable to the site-specific variation exhibited across replicate experiments using the same algorithm (Fig S6).

We also investigated whether the accelerated algorithm had an effect on how epistatic mutations accumulate. A number of recent studies have focused on understanding the influence of epistasis at the level of position-specific substitutions along a protein sequence [6,13,19,31]. A core concept in these works is that substitutions which were neutral or nearly neutral at the time of fixation become entrenched over time, and the probability of reverting back to the substitution’s predecessor diminishes as additional mutations accumulate [19]. We examined the probability of reverting to an ancestral state to assess if our accelerated algorithm caused altered fixation patterns of epistatic mutations during equilibrium behavior. For every substitution, we measured, after each of the subsequent 15 substitutions, the *ΔG* of the protein with the initial mutation reversed and then calculated the probability of fixation of the reversion mutation in the current background. Since the original and accelerated algorithm produced acceptance probabilities on different orders of magnitude, we normalized these probabilities to the sum over the 15 Markov steps, to allow for comparison. Under both algorithms, as substitutions accumulate the probability of accepting a reversion mutation to the ancestral state declines (Fig S7), as expected from the recent literature [6,19]. The rate of decline seems to be slightly faster in the accelerated algorithm, but overall both algorithms show very similar behavior.

### Analysis of selection coefficients and rescaling of *N*_*e*_

We also compared the distribution of fixed selection coefficients during equilibrium behavior (Fig 4). For both algorithms, these distributions have means close to zero, but the distributions themselves are significantly different (KS test; *p* = 4.2 × 10^−3^). Notably, the accelerated algorithm is more conservative and fixes more mutations with smaller selection coefficients. The bulk of mutations fixed by either algorithm are neutral or nearly neutral (defined as |*s*| ≤ 1/*N*_*e*_ [32]), 95% under the original algorithm and 97% under the accelerated algorithm.

**Figure 4.**
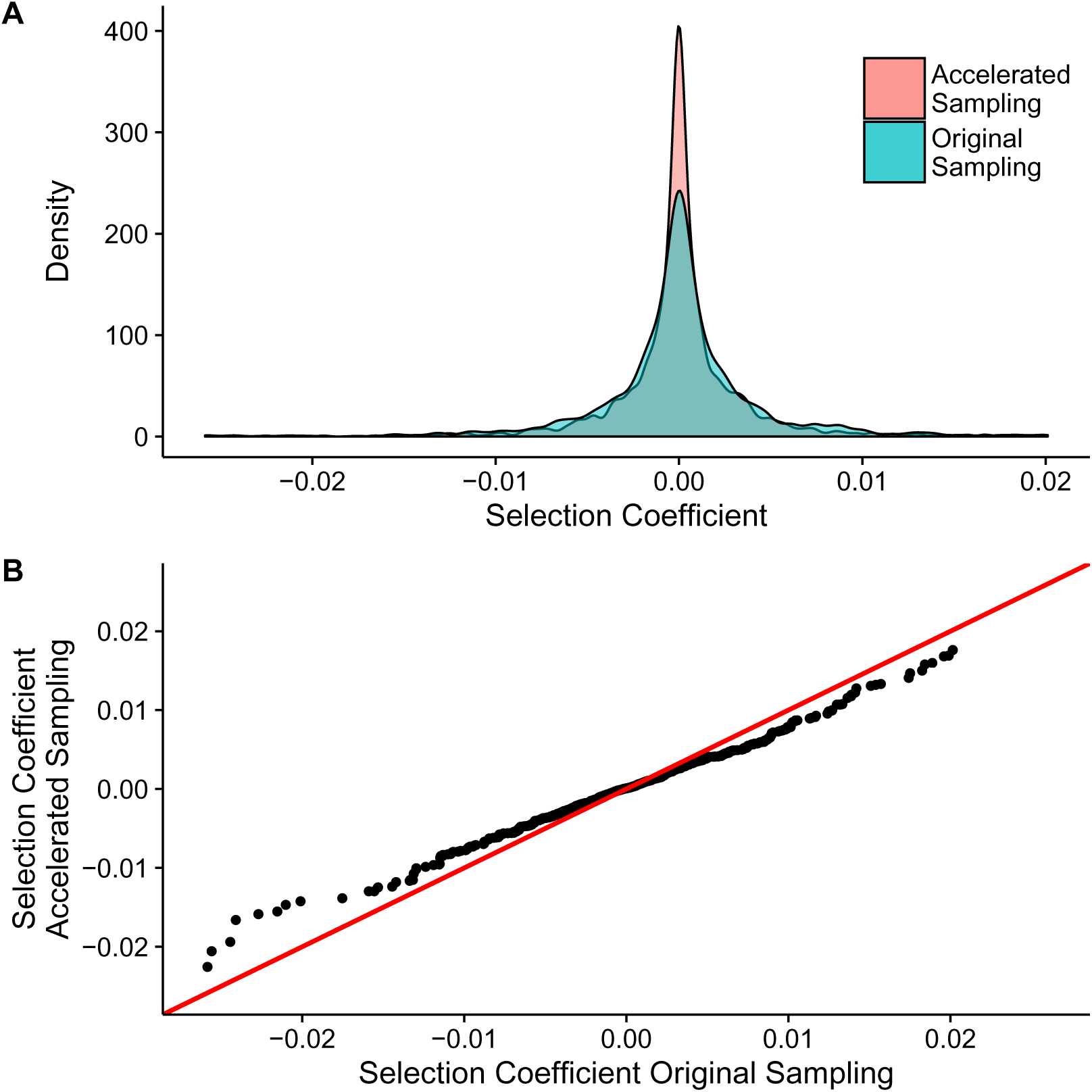
Comparison of the distribution of fixed selection coefficients generated under the original or accelerated sampling. Simulation parameters were as in Fig. 3, but measurements were taken over 500 accepted mutations after an initial burn-in phase of 1000 accepted mutations. (A) Distributions of fixed selection coefficients. The two distributions differ significantly (KS test; *p* = 4.2 × 10^−3^). (B) Q-Q plot of the two distributions of fixed selection coefficients. The *x* = *y* line is given in red. The Q-Q plot highlights systematic differences between the two distributions.

We found that the larger amount of variation in selection coefficients observed in the original algorithm could be recaptured by decreasing *N*_*e*_ in the accelerated algorithm (Fig S8). Specifically, we could rescale *N*_*e*_ to 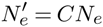, where we chose the constant *C* such that it maximized the *p*-value in a KS test of the two distributions of selection coefficients. We then repeated the accelerated simulations using the rescaled 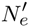 and then compared the distributions of selection coefficients between the original and rescaled accelerated simulations. We found that now the KS test indicated no difference between the two distributions (*p* = 0.72, Fig 5A).

**Figure 5.**
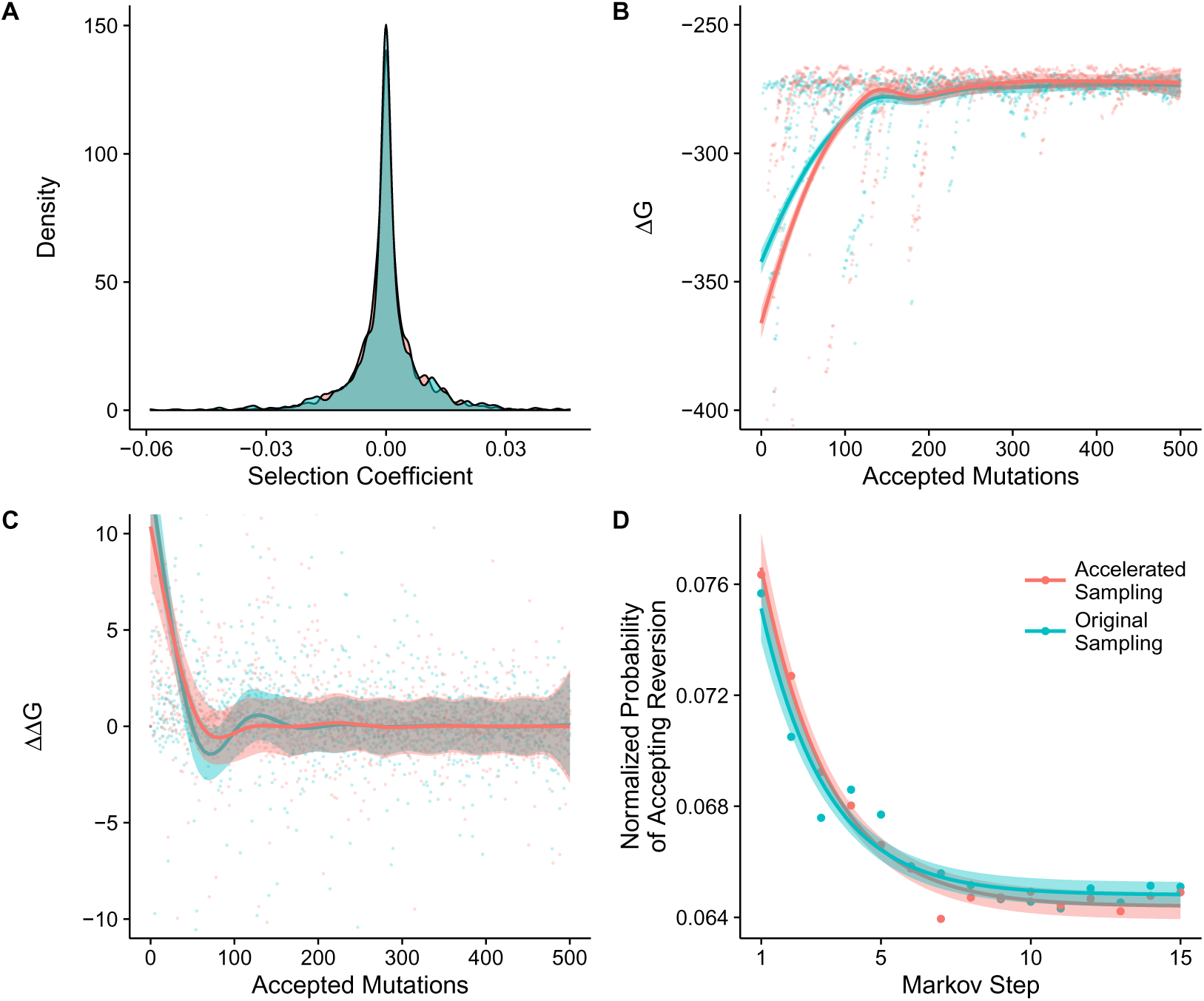
Rescaling *N*_*e*_ in the accelerated algorithm minimizes differences between the two algorithms. Simulation parameters were *β* = 1, *ΔG*_thresh_ = −267, and *N*_*e*_ = 50 or *N*_*e*_ = 37 for the original or the accelerated algorithm, respectively. (A) Distribution of fixed selection coefficients using both algorithms. These distributions are equivalent (KS test; *p* = 0:72). (B) *ΔG* of first 500 accepted mutations. (C) *ΔΔG* of first 500 accepted mutations. (D) Probability of accepting a reversion substitution. Probabilities are normalized to one for each original substitution analyzed. Individual points show the probability averaged over 1500 substitutions (500 substitutions in 3 replicate experiments), the solid line represents a model of exponential decay (*y* = *ce^−ax^* + *b*) fitted to the data (see Table S3 for fitted parameter values), and the shaded area represents the standard error on the fitted curve.

To examine more systematically the effect of rescaling *N*_*e*_ in the accelerated algorithm, we repeated several of the previously outlined analyses after rescaling. We found that rescaling eliminated most of the previously observed differences between the original and the accelerated algorithm (Fig 5B–D). The equilibrium protein stability decreased slightly, though by a barely noticeable amount, when using the accelerated algorithm with rescaled *N*_*e*_ (Fig 5B). However, the amount of variation observed in *ΔΔG* (Fig 5C) was more similar to the original algorithm. The pattern of decay in the probability of accepting a reversion mutation was also now more similar to the original algorithm (Fig 5D).

We determined the optimal scaling constant *C* at two additional values of *N*_*e*_, and we found a linear relationship between the rescaled *N*_*e*_ used in the accelerated algorithm and the *N*_*e*_ used in the original algorithm (Fig 6). This linear relationship suggested that the accelerated algorithm requires an approximately two-times smaller effective population size to generate comparable genetic drift relative to the original algorithm.

**Figure 6.**
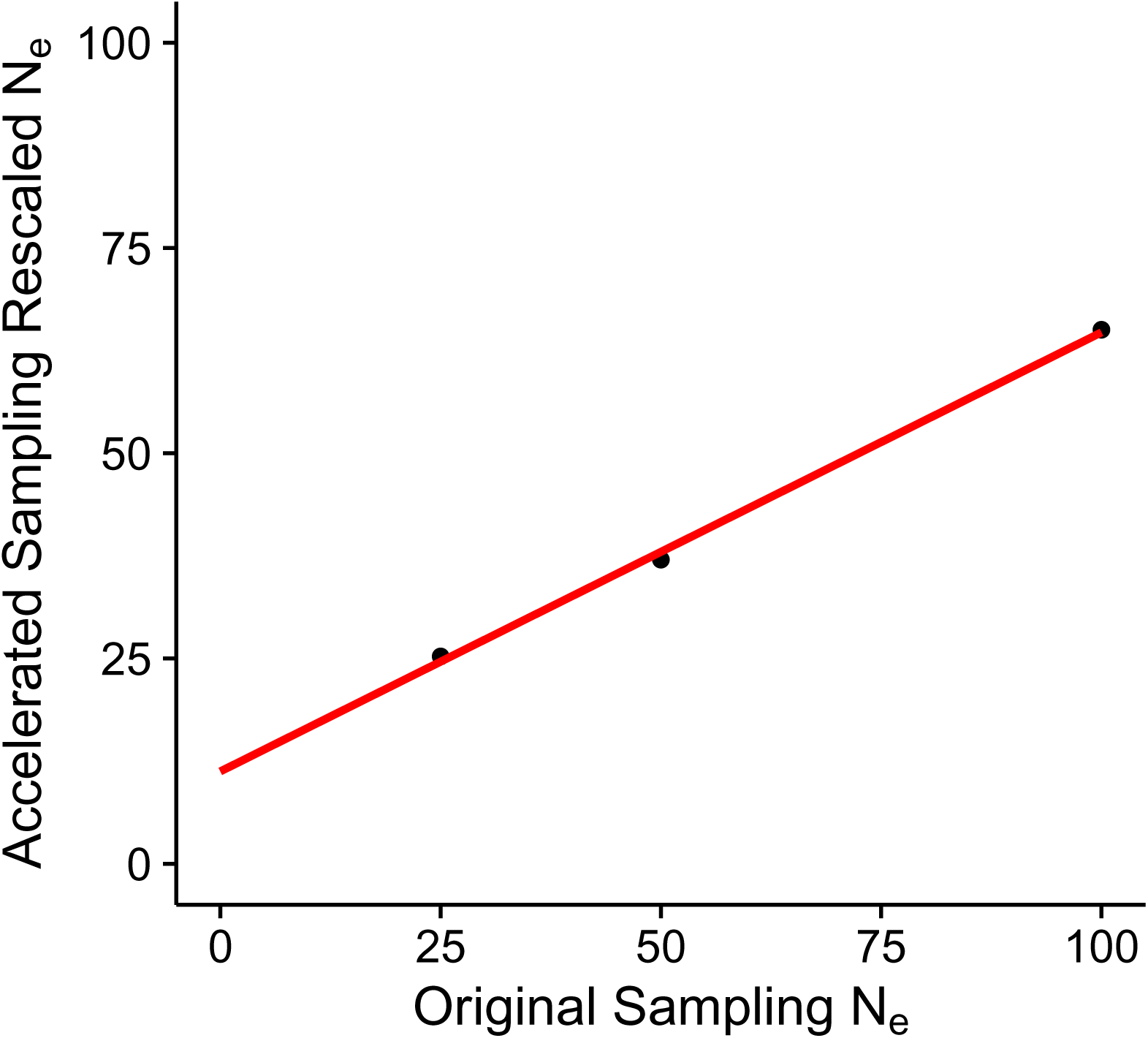
Relationship between the *N*_*e*_ used in the original algorithm and the rescaled *N*_*e*_ required by the accelerated algorithm to produce similar distributions of fixed selection coefficients. Simulation parameters were *β* = 1 and *ΔG*_thresh_ = −267. Simulations using the rescaled *N*_*e*_ in the accelerated algorithm were found to have equivalent distributions of selection coefficients to their original sampling simulation counterparts (KS tests; *p* = 0.75, 0.72, 0.19 for *N*_*e*_ = 100, 50, 25, respectively). The red line represents a linear fit with slope *m* = 0.53 and intercept *b* = 11.25.

## 3. Discussion

We have presented an accelerated algorithm to simulate origin-fixation models. Our algorithm improves run-time efficiency of simulations by approximately an order of *N*_*e*_. For example, in simulations with *N*_*e*_ = 100 we found a 64-fold speed increase relative to the regular algorithm. Our accelerated algorithm does not change the frequency with which states are sampled, but it does alter some aspects of the evolutionary process. For the example case of proteins evolving under a soft threshold model, we have found that our algorithm does not substantially modify equilibrium protein stability, the distribution of accumulated substitutions, or epistatic interactions. However, we have observed that the variance in selection coefficients of fixed mutations is reduced.

Notably, the reduction in variance is relatively minor, and we can recover the original variance by using a slightly reduced population size when simulating with the accelerated algorithm. The scaling between the original and reduced population size is approximately a factor of 2, on the order of the differences between haploid and diploid models or between Wright-Fisher sampling and the Moran process [33]. Of course, reducing N_e_ results in a slightly modified steady state, and for the soft-threshold model most notably a very minor decrease in equilibrium stability, as expected by the intrinsic relationship between population size and protein stability [30]. However, otherwise we have observed little difference between the accelerated algorithm with rescaled population size and the original algorithm with original population size.

While our algorithm demonstrates a considerable performance gain, it also exhibits several shortcomings. First, as stated, our algorithm results in a reduction in the amount of variance sampled along evolutionary trajectories, and thus implicitly simulates a larger effective population size *N*_*e*_ than is used as parameter value in the simulation. However, the implicit increase in *N*_*e*_ is moderate, on the order of a factor of 2. Second, we have emphasized equilibrium dynamics here. Considering that the paths sampled by the two algorithms differ, we expect that the approach to equilibrium will differ between the algorithms. Hence, the two algorithms are likely not going to perform similarly away from equilibrium. Third, we have only compared the behavior of these algorithms in simulations with relatively small values of *N*_*e*_ (*N*_*e*_ ≤ 100). While our algorithm is not expected to perform substantially differently at larger values of *N*_*e*_, we cannot test this hypothesis because of computational constraints associated with using large *N*_*e*_ in the original algorithm. Despite the various shortcomings of our algorithm, for questions that only deal with equilibrium behavior and whose answer doesn’t depend on sampling order or variance in the selection coefficients, the performance gain from using the accelerated algorithm is substantial.

It is well known in population genetics that effective population size, genetic drift, and evolutionary time are closely interconnected. In fact, there are multiple ways to define effective population size based on the aspects of genetic drift in question [34,35]. Considering that our rescaling of the probability of transitions between genotypes is analogous to rescaling time, downstream effects on how genetic drift and *N*_*e*_ operate within our model are to be expected. Examining the reduction of variance in the distribution of selection coefficients and the reduction of variance in states sampled on a simplified landscape, we infer that under our accelerated algorithm drift is more limited than under regular sampling. Considering the inherent relationship between population size and drift, it then follows that we are able to recapture the breadth of drift by decreasing the *N*_*e*_ used in our accelerated algorithm. While the exact linear relationship we found here between the *N*_*e*_ in the original simulation and the *N*_*e*_ in the accelerated simulation may depend on aspects of the particular protein we studied, we expect that similar rescalings will be possible for most fitness landscapes.

The accelerated algorithm we have presented here has been used previously in a few studies [17,36], but without consideration of whether it actually represented an accurate approximation to a realistic evolutionary model. We have shown here that this algorithm may indeed be suitable in a variety of different modeling contexts. The enhanced performance of our algorithm opens the door for a wealth of studies which could benefit from the use of more sophisticated models of protein structure or of other aspects of the fitness landscape. Further applications of our work include examining the role and source of entrenchment with a more rigorous treatment of protein structure than is currently being employed [13,19,20,31]. Our algorithm can also be applied to more detailed studies of the way in which proteins maintain interacting partners over long evolutionary periods [36]. Finally, our algorithm is flexible enough to allow for any fitness function to be used in place of the soft-threshold stability-based model we used here. For example, efficient versions of evolutionary simulations of pathway flux [37] could be constructed and the probability of reversion to an ancestral state could be examined to study the role of epistatic effects at the pathway level.

The field of molecular evolution has long sought to understand the relationship between protein structure and sequence evolution. Here, we have introduced an accelerated simulation algorithm that may serve as a useful tool in pursuing this goal. Our work demonstrates that simplified or coarse-grained models of protein structure and function are not required for simulation studies of protein evolution. Our algorithm is sufficiently fast that fitness landscapes based on full-atom approximations using empirical potentials [38] or composite statistical and physics based potentials [26] can easily be simulated on modern desktop computers. Even though our algorithm does perturb some aspects of the evolutionary process, these perturbations are minor and/or do not influence the main quantities of interest. Thus, our algorithm paves the ground for future simulation studies that use fitness landscapes with much increased levels of realism.

## 4. Methods

### Simulations over a three-state fitness landscape

Simulations over the three-state fitness landscape were performed using custom Python scripts. States A and B were assigned the same fitness (1.0), and state C was given a higher fitness (1.25). *N*_*e*_ = 10 was chosen to allow for adequate sampling of transitions to lower fitness states. At each step in the simulation a random state excluding the state currently occupied was proposed. The proposed state was accepted or rejected based on either Eq. (2.1) (regular algorithm) or Eq. (2.7) (accelerated algorithm). Fifty replicates of each experiment were performed. Each simulation was run for 100,000 iterations, and the states visited and the types of transitions accepted were stored over the last 2,500 iterations. The mean and variance exhibited by the two different acceptance criteria was evaluated by comparing the results from these replicated experiments. The results of these comparisons is given in Fig 2, Table S1, and Table S2.

A similar analysis was carried out on a different fitness landscape were the three states were given fitness values 1.0 (state A), 1.15 (state B), and 1.25 (state C). Data were analyzed as before. The results of these experiments are given in Fig S3, Table S1, and Table S2.

### Simulations of protein evolution

We simulated protein evolution using the all-atom energy function implemented in the Rosetta protein-design suite [25,26]. We implemented both the original and the accelerated algorithm in Python, and we used PyRosetta [39] to mutate the protein structure and to evaluate its energy. We used the Rosetta score as a substitute for *ΔG* in Eq. (2.8). Throughout all calculations, we held the protein backbone fixed.

The protein used for all simulations was a purple acid phosphatase (PDB identifier 1QHW [40]). We first cleaned the structure using the `cleanAtom` function in PyRosetta [39] to remove extraneous information from the PDB file. Then, before each simulation, we minimized the Rosetta score with the Davidon-Fletcher-Powell gradient minimization method provided through PyRosetta [39] (function `dfpmin_armijo_nonmonotone`). Starting from the minimized structure, either simulation algorithm then proceeded as follows. At each discrete time step, we randomly chose a site in the structure and replaced the resident amino acid with one of 18 alternative amino acids, also chosen at random. The alteration of cysteine residues present in the initial structure and the addition of cysteines through mutation was not allowed to avoid modifying or adding disulfide bonds. We then locally repacked the structure 10 Ångstroms around the mutation and calculated the Rosetta score. Finally, we used either Eq.(2.1) (for the original algorithm) or Eq.(2.7) (for the accelerated algorithm) to determine whether to accept or to reject the mutation.

We set *N*_*e*_ = 100 to keep simulations computationally tractable when using the original algorithm. The parameter *β* was set equal to 1, and *ΔG*_thresh_ was set to half of the Rosetta score of the initial structure, which was −534, measured in arbitrary units. To assess the steady state dynamics of the two algorithms we ran each trajectory in triplicate until we had accumulated 1,500 substitutions. To insure that steady state sampling behavior was being exhibited we discarded the first 1,000 substitutions as burn-in before preforming analysis comparing steady state behavior. For the remaining 500 substitutions, we recorded the number of substitutions observed at each position in the protein, the probability of accepting a reversion mutation, and the selection coefficients of fixed mutations. To examine if the two algorithms differed in their initial evolutionary behavior as they approached equilibrium we carried out identical simulations to the ones described above but recorded only the first 500 substitutions.

The probability of accepting a reversion mutation was calculated using data from simulations run for 1,500 substitutions where the first 1,000 substitutions were discarded as burn-in. The probability of accepting a reversion mutation was determined by calculating the fitness of the ancestral state at each of the following 15 Markov steps after the acceptance of every new state, and then evaluating the probability of acceptance of the reversion using either model. This data was arranged in a matrix with 15 rows, one for each Markov step, and 1,500 columns, one for each substitution (500 substitutions for each of the 3 replicates). Each column in the matrix was normalized such that the probability of any specific substitution being accepted along the following 15 Markov steps summed to 1. The row means were then computed from this column normalized matrix to give the mean probability of accepting a reversion mutation at any specific Markov step.

We carried out additional simulations to test the rescaling of the effective population size of 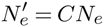. First, we used the original sampling to simulate trajectories with 1,500 substitutions at population sizes of *N*_*e*_ = 25, 50, 100, each performed in triplicate. We then simulated trajectories with the same *N*_*e*_ using accelerated sampling. Next, for each population size, we chose an appropriate scaling constant *C* that maximized the *p*-value in a KS test of the distributions of selection coefficients obtained under original and accelerated sampling. Finally, we simulated trajectories with the corresponding 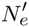 values, again performed in triplicate.

### Statistical analysis and data availability

Statistical analysis was performed with R [41]. Data visualization and model fitting were performed with ggplot [42]. All source code and processed data are available at https://github.com/a-teufel/Accelerated_Sim.

## Acknowledgments

We thank Austin Meyer and Julian Echave for useful discussions.

## Author Contributions

COW conceived of the project. Experiments were designed by AIT and COW and performed by AIT. The manuscript was written by AIT and COW.

